# Comprehensive Genetic Testing for Female and Male Infertility UsingNext Generation Sequencing

**DOI:** 10.1101/272419

**Authors:** Bonny Patel, Sasha Parets, Matthew Akana, Gregory Kellogg, Michael Jansen, Chihyu Chang, Ying Cai, Rebecca Fox, Mohammad Niknazar, Roman Shraga, Colby Hunter, Andrew Pollock, Robert Wisotzkey, Malgorzata Jaremko, Alex Bisignano, Oscar Puig

## Abstract

**Objective:** To develop a comprehensive genetic test for female and male infertility in support of medical decisions during assisted reproductive technology (ART) protocols.

**Design:** Retrospective analysis of results from 118 DNA samples with known variants in loci representative of female and male infertility.

**Interventions(s):** None

**Main Outcome Measure(s):** Next-Generation Sequencing (NGS) of 87 genes including promoters, 5’ and 3’ untranslated regions, exons and selected introns. In addition, sex chromosome aneuploidies and Y chromosome microdeletions are analyzed concomitantly using the same panel.

**Results:** Analytical accuracy was >99%, with >98% sensitivity for Single Nucleotide Variants (SNVs) and >91% sensitivity for insertions/deletions (indels). Clinical sensitivity was assessed with samples containing variants representative of male and female infertility, and it was 100% for SNVs/indels, *CFTR* IVS8-5T variants, sex chromosome aneuploidies and Copy Number Variants (CNVs), and >93% for Y chromosome microdeletions. Cost analysis comparing the NGS assay with standard, multiple analysis approach, shows potential savings of $2723 per case. Conclusion: A single, comprehensive, NGS panel can simplify the ordering process for healthcare providers, reduce turnaround time, and lower the overall cost of testing for genetic assessment of infertility in females and males, while maintaining accuracy.

## Introduction

It is estimated that 48 million couples were affected by infertility in 2010, and there has not been any significant improvement on infertility levels between 1990 and 2010 (1–3). In the United States alone, 12% of women aged 15-44 have impaired fecundity. More couples are relying on assisted reproductive technologies (ART) to get pregnant and have children, and over 1.6 million ART cycles were performed worldwide in 2010 (4). The inability to have children affects couples worldwide and causes emotional and psychological distress in both men and women.

It has been estimated that a very high proportion of infertility cases are due to genetic defects. Male infertility accounts for 50% of the cases (5), with known genetic factors being responsible for 15-30% of male infertility (6). Chromosomal alterations (7), inversions (8), translocations (9), Y chromosome microdeletions (10) and gene mutations (for example Single Nucleotide Variants (SNVs) in *CFTR* (11)) are the main genetic variants causing male infertility. In females, infertility is more of a heterogeneous condition where genetics clearly play a role, but mostly with polygenic effects making it difficult to define a single genetic cause. Exceptions include sex chromosome alterations (12) and several single gene mutations (13,14), but the overall prevalence of these seem to be low (15).

Fertility treatments are costly (∼$12,400/cycle) and effective only in a subset of patients (overall success rate for live birth is 29%)(16). As a consequence, couples frequently undergo several In Vitro Fertilization (IVF) cycles, a process spanning several months and with median out-of-pocket costs in the USA over $60,000 (17). Unfortunately, success rates are not predictable, and some couples discontinue before achieving pregnancy due to economical or psychological factors (18), while others continue despite the presence of an identifiable genetic defect that will prevent success. For couples who will ultimately be unable to conceive even through IVF, a fast, accurate and inexpensive diagnostic test that identifies causes of infertility would be very useful to help them seek alternative solutions early in the process (e.g. adoption or sperm/egg donors). Those in whom genetic defects are not identified might be encouraged to undertake or continue IVF until success is achieved.

Traditionally, several assays are needed to make a definitive genetic diagnosis of infertility, which makes the process expensive and slow. For example, in males, sex chromosome aneuploidies are detected by cytogenetic tests like karyotyping, Y chromosome microdeletions are detected by Polymerase Chain Reaction (PCR)-based methods, and *CFTR* mutations by Sanger DNA sequencing. However, a shotgun approach in which all genetic tests are ordered for all patients is not recommended because cost is prohibitive (19).

Next generation sequencing (NGS) permits the simultaneous interrogation of multiple disease-causing variants in many genes, allowing expanded genetic diagnostic to be routinely used in medical practice. This is already a reality in other medical fields, like oncology (20) and heart disease (21), where panel testing allows for the most comprehensive assessment of genetic etiologies. NGS is also very cost-effective because it allows for the detection of very different types of variants (for example, SNVs, small indels, large Y chromosome deletions and sex chromosome aneuploidies) by using a single test in combination with multiple bioinformatics algorithms to process these diverse data. Here, we present the development of an NGS panel and bioinformatics pipeline for the detection of genetic variants with direct impact in female and male infertility. We also present a cost comparative analysis of current approaches in comparison with the NGS test.

## Materials and Methods

### Subjects and Samples

Samples of genomic DNA with previously identified variants in the genes interrogated by the sequencing panel were obtained as purified DNA from the Human Genetic Cell Repository (National Institute of General Medical Sciences) at the Coriell Institute for Medical Research (Camden, NJ) or the Indiana Biobank (Indianapolis, IN), or extracted from saliva samples collected by DNAsimple (Philadelphia, PA). All samples submitted to the Phosphorus Diagnostics laboratory as saliva were manually extracted using the Qiagen QIAamp Mini Kits following the manufacturer’s instructions. DNA was quantified using Quant-iT Picogreen dsDNA Assay Kit (Life Sciences) and a Varioskan LUX (Thermo Scientific). Supplementary Table S1 describes all samples used in this study, with biorepository numbers, as well as their previous characterization results and rationale for inclusion. All samples presented here are de-identified (HHS 45 CFR part 46.101(b)(4)). IRB approval to handle de-identified samples was obtained through ASPIRE (Santee, CA), protocol IRB-R-003.

### Fertility panel description

A targeted next generation sequencing panel consisting of 87 genes related to infertility disorders was created (Supplementary Table S2). Genes relevant to disease phenotype were included based on relationships described in Online Mendelian Inheritance in Man (OMIM), Human Phenotype Ontology (HPO), GeneReviews, and primary literature. The panel included all coding exons, splice sites, promoter regions, 5’-Untranslated Regions (UTRs), and 3’-UTRs for each of the genes. Clinically relevant noncoding (intronic) regions that contained previously described pathogenic variants, as reported in ClinVar NIH database, were also included. The total size of the gene panel is 1,444,982 bp. In addition, selected regions from the Y chromosome (928,649 bp) were included to allow the detection of Y chromosome microdeletions.

### Next Generation DNA Sequencing

All laboratory procedures were performed in a Clinical Laboratory Improvement Amendments (CLIA) laboratory (Phosphorus, New Jersey, NJ). DNA samples were prepared for sequencing using HyperPlus Library Preparation Kit (Roche, Indianapolis, IN) and sequenced on a NextSeq500 (Illumina, San Diego, CA), following the manufacturer’s instructions. A detailed sequencing protocol is provided in Supplementary Methods section.

### Variant identification and classification

All bioinformatics algorithms were implemented within the Elements^™^ platform (Phosphorus, New York, NY). FASTQ files were produced from each sequencing run and processed using the germline calling pipeline (version 2.03.01.30066) in DRAGEN (Edico Genome, San Diego, CA). Variants identified by NGS were confirmed by an orthogonal method (microarrays or Sanger sequencing). After confirmation, each variant was classified as pathogenic, likely pathogenic, variant of unknown significance (VUS), likely benign or benign, following the American College of Medical Genetics (ACMG) guidelines (22).

### Orthogonal confirmation

A custom Phosphorus Affymetrix axiom array was used to confirm all the reportable variants including SNVs, indels, CNVs, Y chromosome microdeletions and sex chromosome aneuploidies. Microarrays were processed following the manufacturer’s instructions. To confirm variants not included in the microarray, Sanger sequencing was used. Orthogonal analysis was also performed for cases when there was discrepancy between expected and received NGS results. If both NGS and confirmatory results agreed, the results were counted as concordant. Detailed protocols are described in the Supplementary Methods section.

### FMR1 testing

*FMR1* testing was performed with 120 ng genomic DNA per sample and the AmplideX PCR CE *FMR1* kit (Asuragen, Austin, TX), following the manufacturer’s instructions. PCR products were separated in a 3500XL capillary electrophoresis system (Applied Biosystems, Foster City, CA) using conditions described in the *FMR1* kit manual.

## Results

### Gene Panel Design

In order to develop a comprehensive infertility genetic test, we focused only on genes with demonstrated impact on infertility phenotype. Genes were classified as “diagnostic” when variants in them are described as causing infertility across multiple populations, as supported by multiple publications from different laboratories, and showing a direct relationship with infertility. Genes were classified as “informative” when variants in them have been described to be associated with infertility, but the causality link has not been unequivocally established. For male infertility, the panel included Y chromosome microdeletions, *CFTR* mutations and sex chromosome aneuploidies (6). For female infertility, the panel included sex chromosome aneuploidies and genes in which variants have been associated with recurrent pregnancy loss caused by thrombophilia, primary ovarian insufficiency, polycystic ovary syndrome and ovarian hyperstimulation syndrome. The gene list, as well as the rationale used for their selection is shown in Supplementary Table S2.

### Analytical and Clinical Validation

In order to validate panel performance and determine analytical sensitivity, specificity and accuracy, we used 24 samples from the 1000 Genomes (1000G) project (23) for which the location of SNVs and indels are known (Supplementary Table S1). Representative sequencing quality control statistics are shown in Supplementary Table S3. NGS sequencing results from these validation samples was compared known 1000G variants (Table 1). Microarray analysis and/or Sanger sequencing was performed to assess the discrepancies of SNVs/indels between results produced by the NGS panel and previously known 1000G data. Analytical sensitivity of the test was >99% for SNVs and >91% for indels, and specificity was >99% for both SNVs and indels. Final accuracy for SNVs and indels was 99.98% and 99.42%, respectively.

**Table 1.**
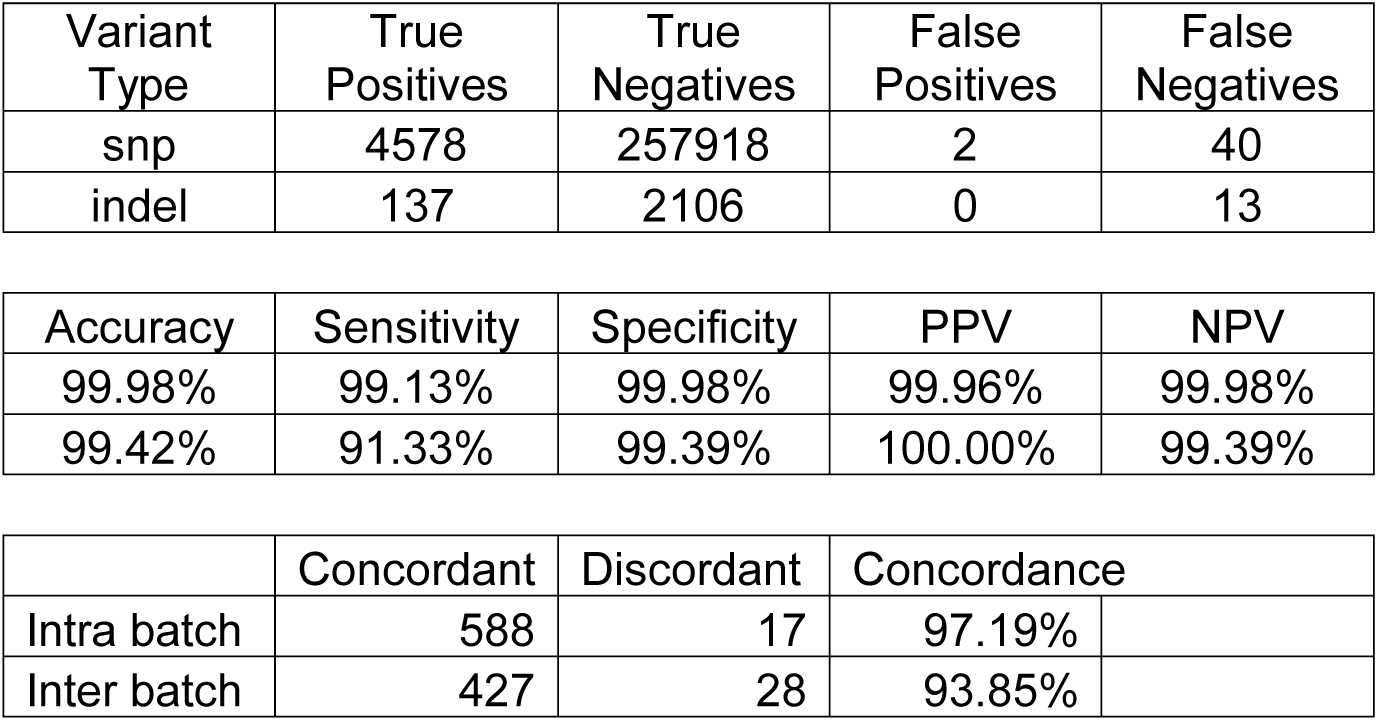
Performance characteristics of the NGS test. PPV, Positive Predictive Value. NPV, Negative Predictive Value. CNV, Copy Number Variant

To determine clinical sensitivity in the detection of CNVs, sex chromosome aneuploidies and Y chromosome microdeletions, we used 34 samples with 38 known variants (Supplementary Table S1). The set of variants previously known for these samples is limited, so specificity could not be assessed. The analysis correctly detected 3/3 CNVs, 19/19 sex chromosome aneuploidies and 15/16 Y chromosome microdeletions (Supplementary Tables S4-S5, Table 1 and Figure 1). In three cases (NA20435, NA18333 and NA22031) Y chromosome microdeletions of smaller size were found by NGS when compared to previously known data (Supplementary Table S5), however, microarray analysis confirmed the previously reported microdeletion size. In one case (NA20434), NGS data located the microdeletion 1.17 Mbp downstream of the previously known location, and microarray analysis confirmed NGS result. Sample NA12662 has a sex chromosome aneuploidy (duplication of X chromosome) and a Y chromosome microdeletion ((22769319-27097245)x0). The sex chromosome aneuploidy was identified, but the Y chromosome microdeletion was missed. Thus, clinical sensitivity for sex chromosome aneuploidies and CNVs is 100% (22/22), and for Y chromosome microdeletions is 93.75% (15/16).

**Figure 1.**
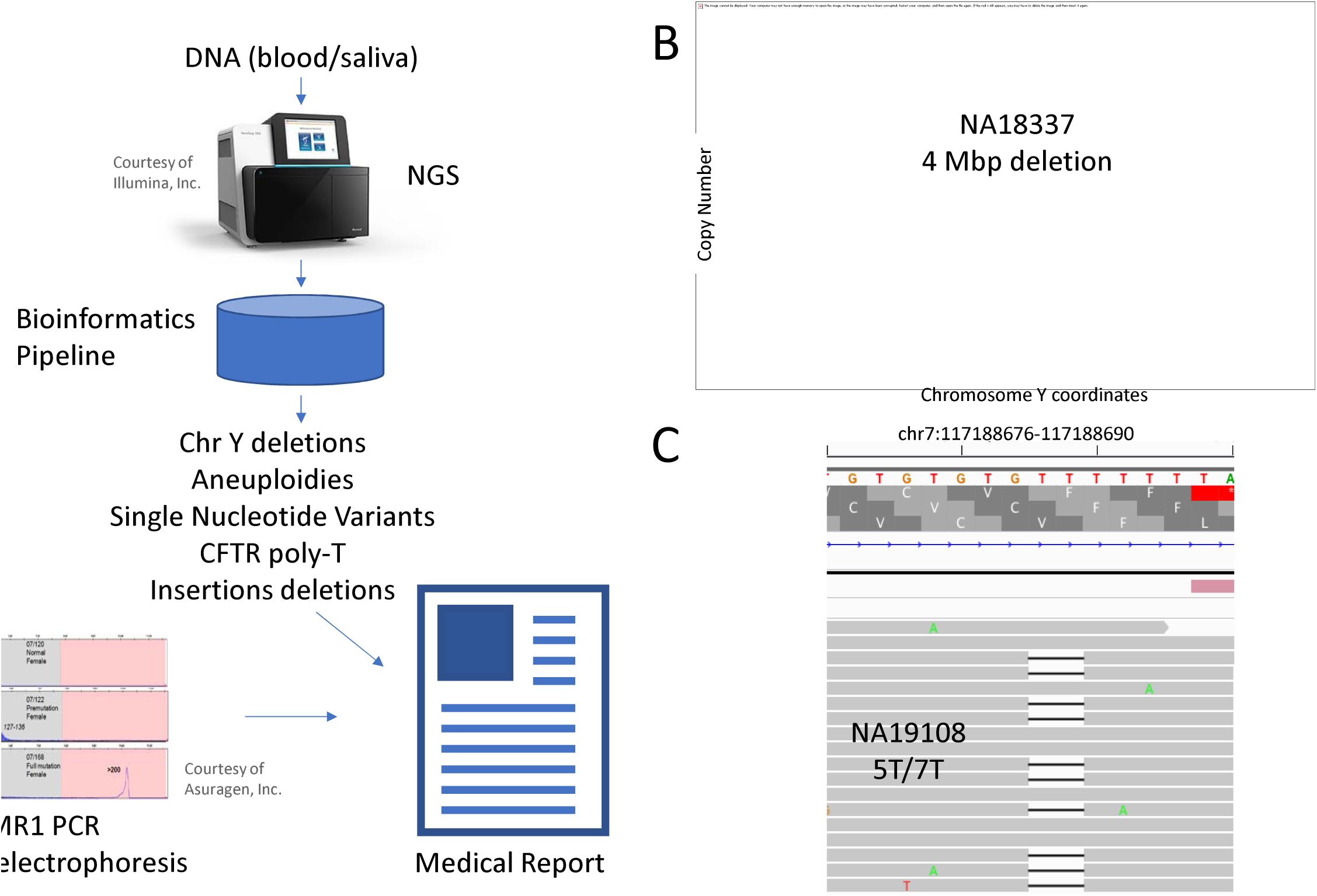
A) Outline of our NGS test. A DNA sample (saliva or blood), is sequenced by NGS and processed by our custom bioinformatics pipeline. Y Chromosome microdeletions, sex chromosome aneuploidies, *CFTR* IVS8-5T polymorphism, indels and SNVs are called. Variants are interpreted by expert curators and a medical report is generated. In parallel, *FMR1* testing is performed using PCR and capillary electrophoresis, and results are incorporated into the medical report. B) Example of Y chromosome microdeletion called in sample NA18337 C) Example of IVS8-5T tract detection in sample NA19108, heterozygous for 5T/7T.

Next, 11 DNA samples with 17 known SNV/indels were processed (Table 2). Known variants were confirmed in 16/17 cases. Sample NA02795 was reported with a pathogenic variant *GALT* c.130G>A; p.Val44Met. However, this variant was not identified by NGS and Sanger sequencing did not detect it either, therefore, based on NGS and Sanger sequencing concordance, we did not count this case against the sensitivity. Thus, clinical sensitivity for SNVs/indels by our assay was 100%.

**Table 2.** Clinical sensitivity was determined by sequencing SNVs/indels of known variants

| Sample | Previously reported | NGS test call |
| --- | --- | --- |
| NA11763 | *GALT* c.591A>G; p.Gln188Arg | *GALT* NM_000155.3:c.563A>G; p.Gln188Arg |
| NA11763 | *GALT* c.1025C>T; p.Arg333Trp | *GALT* NM_000155.3:c.997C>T; p.Arg333Trp |
| NA14899 | *F5* c.1691G>A; p.Arg506Gln (homozygous) | *F5* NM_000130.4:c.1691G>A; p.Arg506Gln |
| IndianaBiobank 02 | *SERPINC1* c.1148_1149delTC; p.Leu383Profs*10 | *SERPINC1* NM_000488.3:c.1148_1149delTC; p.Leu383Profs*10 |
| NA00422 | *GALT* c.563A>G; p.Gln188Arg | *GALT* NM_000155.3:c.563A>G; p.Gln188Arg |
| NA00422 | *GALT* c.1030C>A; p.Gln344Lys | *GALT* NM_000155.3:c.1030C>A; p.Gln344Lys |
| NA02795 | *GALT* c.425T>A; p.Met142Lys | *GALT* NM_000155.3:c.425T>A; p.Met142Lys |
| NA02795 | *GALT* c.130G>A; p.Val44Met | None |
| DNAsimple01 | *CFTR* c.1584G>A; p.Glu528= | *CFTR* NM_000492.3:c.1584G>A; p.Glu528= |
| NA02796 | *GALT* c.512T>C; p.Phe171Ser | *GALT* c.512T>C; p.Phe171Ser |
| NA02796 | *GALT* c.404C>T; p.Ser135Leu | *GALT* c.404C>T; p.Ser135Leu |
| NA16643* | *F5* c.1691G>A; p.Arg506Gln | *F5* c.1601G>A; p.Arg534Gln |
| NA17431 | *GALT* c.591A>G; p.Gln188Arg | *GALT* NM_000155.3:c.563A>G; p.Gln188Arg |
| NA17431 | *GALT* c.612T>C; p.Leu195Pro | *GALT* NM_000155.3:c.584T>C; p.Leu195Pro |
| NA16000 | *MTHFR* c.677C>T; p.Ala222Val | *MTHFR* c.677C>T; p.Ala222Val |
| NA16000 | *F2* c.20210G>A (homozygous) | *F2* NM_000506.4:c.*97G>A |
| DNAsimple02 | *F2* c.20210G>A | *F2* NM_000506.4:c.*97G>A |

\*The official designation of *F5* c.1691G>A is NM\_000130.4:c.1601G>A; NP\_000121.2:p.Arg534Gln

### CFTR intron 8 poly dT (IVS8-5T)

Varied lengths of a thymidine (T)-tract (5, 7, or 9T) are found in front of the splice-acceptor site of intron 8 of the *CFTR* gene in males with congenital bilateral absence of the vas deferens (24). The length correlates with the efficiency of exon 9 splicing. This polymorphism is detected clinically by allele-specific multiplex PCR (25). In order to determine sensitivity of the NGS panel, we analyzed 72 samples from the 1000G project. When 1000G data was used as reference, the NGS panel correctly detected the IVS8 alleles in 67/72 cases, and for the 5 discrepant cases, Sanger sequencing demonstrated that the NGS panel call was the correct one (Supplementary Table S6). Therefore, our assay had 100% sensitivity to detect the *CFTR* IVS8-5T allele.

### FMR1 testing

CGG triplet expansions are known to cause fragile X syndrome, and alleles in the permutation range have been associated with increased risk for premature ovarian failure (26). Current NGS protocols based on hybridization enrichment for targeted panels do not accurately quantify the number of repeats. Therefore, to complement the NGS panel with *FMR1* analysis we used an already available protocol based on PCR amplification and capillary electrophoresis detection of CGG repeats (27). Test sensitivity was determined by analysis of 26 samples harboring different CGG repeat expansions (Supplementary Table S7). We correctly identified all alleles previously classified as “mutation” (>200 CGG repeats), “pre-mutation” (55 to 199 CGG repeats) and “normal” (<45 CGG repeats). However, the assay misclassified one allele in the “intermediate” range (between 45 and 54 CGG repeats) as “pre-mutation” (NA20236, 53 vs 55 CGG repeats, Supplementary Table S7). Given that the rate of expansions of intermediate alleles is not well understood, testing at-risk relatives of individuals with an intermediate allele may determine the stability of the allele in the family (28).

### Cost analysis

Currently, genetic testing for variants associated with male infertility is performed with multiple assays. For example, *CFTR* IVS8-5T is detected by allele specific multiplex PCR (25), Y chromosome microdeletions are detected by PCR of sequence tagged-sites (29), and sex chromosome aneuploidies are detected by traditional karyotyping or microarrays (30). NGS can detect all these variants using a single test. In order to determine the economic impact, we performed a cost analysis for the male fertility panel, using average pricing available from reference laboratories performing these assays (Table 3 and Supplementary Table S8). Overall, the savings to providers are $2723 per case, $3322 if performed by multiple assays vs $599 if performed by NGS, which translates in savings of >550%. Furthermore, NGS testing has a shorter turnaround time because there is no need to do reflex testing. The only exception may be cases with complex chromosomal rearrangements and mosaics or translocations/inversions that do not result in gain/loss of DNA and are, therefore, not captured by the NGS panel.

**Table 3.** Cost comparison between traditional diagnostic methods for male infertility and Phosphorus’ NGS diagnostic test.

| Variant | Methodology | CPT Code | Cost (\$US) |
| --- | --- | --- | --- |
| CFTR IVS8 poly T tract | Multiplex Polymerase Chain Reaction, sanger sequencing or NGS | 81224 | 396 |
| CFTR mutations | Sanger sequencing of CFTR gene | 81223 | 1430 |
| Sex chromosome aneuploidies | Karyotype | 88262 | 946 |
| Y chromosome microdeletion | PCR-based analysis of 14 different regions along the length of the Y chromosome | 81403 | 660 |
|  |  | Total Cost: | 3322 |
| NGS infertility test | NGS sequencing, includes confirmation with orthogonal methods (microarray or Sanger) | 81224, 81223, 88262, 81403 | 599 |

## Discussion

Genetic analysis of variants linked to infertility is currently performed using many different platforms: SNVs and indels are detected by Sanger sequencing or NGS, sex chromosome aneuploidies by karyotyping or microarrays, Y chromosome microdeletions by multiplex PCR of sequence tags, and *CFTR* intron 8 polymorphism by allele-specific multiplex PCR. NGS allows for the detection of these different variants using a single platform with excellent sensitivity and specificity. Out of 127 different events, the NGS panel complemented with orthogonal confirmation methods accurately detected almost all of them: 17/17 SNVs/indels, 3/3 CNVs, 19/19 sex chromosome aneuploidies, 15/16 Y chromosome microdeletions and 72/72 *CFTR* IVS8 polymorphic sites (5 of them IVS8-5T). In three Y chromosome microdeletion cases, the NGS panel identified microdeletions of smaller size, however, analysis using microarrays confirmed the previously known size in all three cases. The NGS panel, thus, identified accurately most Y chromosome microdeletions (one Y chromosome microdeletion was missed), but orthogonal methods are required to confirm the microdeletion size. In summary, overall clinical sensitivity for all variants is 99.21% (126/127).

Our study has several limitations. Sample size is limited, and even though we processed samples representing all types of variants, the final number of samples in each group is small. For example, we only processed 3 clinical samples containing CNVs. Therefore, given limited sample size, performance characteristics must be interpreted with caution.

The NGS panel cannot detect balanced translocations, therefore it would miss reciprocal translocations and Robertsonian translocations, which are known to cause infertility in 0.9% of men (9). It cannot detect complex chromosomal rearrangements like derivative chromosomes or mosaicism, even though these often involve gain or loss of DNA. We tested several samples with derivative chromosomes and mosaicism and the NGS panel could not accurately assess these variants. For example, samples 3233489 and NA21681 in Supplementary Table S4 contain a derivative X,Y chromosome and a 6,Y translocation, respectively, and both were missed. In these cases, a karyotype is needed, which could be used as reflex after the NGS test. As NGS evolves, it may be possible in the near future to incorporate detection of these abnormalities.

Another limitation is the detection of *FMR1* variant caused by triplet CGG expansion, varying from 20 to >900 repeats. It is not possible to accurately identify triplet expansions by using PCR-enriched NGS. New methods involving PCR-free libraries allow diagnosis (31), but it would require a parallel library preparation and sequencing reaction, significantly increasing cost. Therefore, currently, a separate PCR assay to determine *FMR1* mutation is needed to assess fragile-X mutation.

The direct consequence of using a single NGS assay instead of multiple diagnostic assays, each one detecting a class of DNA variants, is a potential reduction in cost and turnaround time required to make a definitive diagnosis, with savings of $2723 per case ($3322 for traditional methods vs $599 for NGS testing). Integrating testing into a single assay simplifies test ordering and result tracking for the clinician, and decreases cost to the patient by reducing the number of assays that need to be performed, analyzed, and reported.

## Conclusions

We have developed a comprehensive genetic test for female and male infertility that achieves excellent clinical sensitivity, but at a fraction of the cost of traditional methods of testing, and a shorter turnaround time than current methods.

## Authors’ contributions

SP, AB and OP participated in study design and data analysis.

SP, BP, MA, GK, MJ, CC, YC, RF, MN, CH, AP, RW, AB, OP participated in data collection, interpretation of results and drafting the manuscript.

## Acknowledgements

We thank Sara Bristow for her initial scientific input, Santiago Munne for his scientific support and Ed O’Neill for editorial comments on the manuscript.

